# Identification of a specific granular marker of zebrafish eosinophils enables development of new tools for their study

**DOI:** 10.1101/2024.04.29.591640

**Authors:** Miriam Herbert, Christian Goosmann, Volker Brinkmann, Christiane Dimmler, Mark R. Cronan

## Abstract

Eosinophils control many aspects of the vertebrate innate immune response. They contribute to homeostasis, inflammatory conditions and defense against pathogens, yet, their function in disease often remains enigmatic. The zebrafish has emerged as a useful model organism for human diseases but tools to study eosinophils in this model are severely limited. Here, we characterize a new and highly specific marker for eosinophils in zebrafish and report a new transgenic reporter line to visualize eosinophils *in vivo*. In addition, we created a polyclonal antibody that allows the identification of eosinophils *ex vivo*. These new tools expand the toolkit to study eosinophils in the zebrafish model. Using our tools, we have been able to identify the likely ortholog of eosinophil major basic protein, a critical mediator of eosinophil function, in zebrafish. These advances will allow for the investigation of eosinophil biology in the zebrafish model organism, allowing researchers to identify the contribution of eosinophils to the many diseases that are modeled within zebrafish and also shed light on the evolution of eosinophils within vertebrates.

## Introduction

Zebrafish (*Danio rerio*) are an increasingly popular model organism in biomedical research and have been used to model a variety of human diseases^1–3^. They possess an evolutionarily conserved vertebrate immune system containing all major cell types, including eosinophils^4^. The zebrafish’s naturally transparent larval stage, high fecundity and genetic tractability make it an excellent model organism to study hematological and immunological questions^5,6^. While eosinophils are a recognized part of the zebrafish immune repertoire, the limited array of existing tools for the study of eosinophil function in zebrafish has restricted the use of this model for the study of eosinophil biology.

Eosinophils belong to the group of granulocytes and are characterized by large cytoplasmic granules, that store a plethora of pre-formed signaling and effector molecules, which can be stained with acidic dyes like eosin^7^. They are well known effector cells in parasitic infections as well as mediators of allergic or inflammatory conditions, e.g. asthma. Although they are only found in low numbers under homeostatic conditions, they have been associated with both beneficial as well as detrimental effects in health and disease^8–10^. Yet, their contributions often remain elusive.

Past work characterizing eosinophil responses in zebrafish have mostly relied on classical histological staining approaches and the transgenic *Tg(gata2:eGFP)* line^11^. In this transgenic line, the *gata2a* promoter drives the expression of EGFP. Since eosinophils express this marker highly, this transgenic line can be used to isolate eosinophils using fluorescence assisted cell sorting (FACS). However, *gata2a* is not expressed exclusively in eosinophils and therefore this reporter line cannot be used to identify eosinophils *in vivo*.

The broad importance of eosinophils in a range of conditions as well as their evolutionary conservation highlight the critical role played by this cell type and the need for comprehensive methods to further investigate their function in a range of settings. Here, we identify an eosinophil granule-specific marker in zebrafish that we use to create both a transgenic fluorescent reporter line as well as a polyclonal antibody to visualize zebrafish eosinophils both *in vivo* and *ex vivo*. We found that our eosinophil reporter line and antibody could be used to identify eosinophils in tissue as well as by flow cytometry, facilitating the study of this immune cell population. Using electron microscopy, we found that this marker localizes specifically to eosinophil granules. Based on our findings, we are able to identify our marker gene as the zebrafish orthologue of eosinophil major basic protein *(embp)*. To further demonstrate the utility of our tools, we visualized the localization of eosinophils to mycobacterial granulomas, validating the use of this marker during inflammatory conditions.

## Results

### Embp is a granular marker for zebrafish eosinophils

Previous studies into zebrafish eosinophils have often used broadly expressed markers, like the *Tg(gata2:eGFP)* line, or histological properties to identify eosinophils. Single cell RNA-Seq (scRNA-Seq) data allow clustering and annotation of cell types based on transcriptomic profiles. These analyses facilitate an unbiased investigation into uncharacterized gene expression patterns. Using previously published zebrafish scRNA-Seq data sets (GSE161712, GSE112438), we searched for differentially and highly expressed genes in eosinophil populations. One gene that stood out was *dkeyp75b4*.*10* (also known as *eslec*), a previously uncharacterized protein with a C-type lectin domain that is highly and differentially expressed in eosinophils. A recent report established that the promoter of this gene is eosinophil-lineage specific^23^. Based on our findings described below, we have renamed this gene *embp* (for eosinophil major basic protein) which we will use throughout for clarity purposes.

Our aim was to create a transgenic reporter line with a precise knock-in at the endogenous gene locus using CRISPR Cas. By tagging the C-terminus of Embp with a fluorescent protein, this approach will result in the generation of a full-length, tagged version of the protein under the control of the endogenous promoter, allowing us to visualize both eosinophils, and the localization of the Embp protein itself. A donor vector was designed (Fig 1A) to generate this transgenic line. Next, a sgRNA was designed to create a double-strand break with CRISPR Cas9 at the before-last intron of *embp*. The donor DNA fragment reconstitutes the remaining intronic sequence as well as the two last exons downstream of the cut site upon integration. Furthermore, it introduces a tdTomato sequence as a 3’ tag of our gene of interest, followed by a polyA sequence. The construct also includes a transgenesis marker using an eye-specific *cryaa* promoter to drive EGFP enabling facile identification of embryos carrying this transgene prior to the emergence of eosinophils, by bright green fluorescence in the lens.

**Figure 1.**
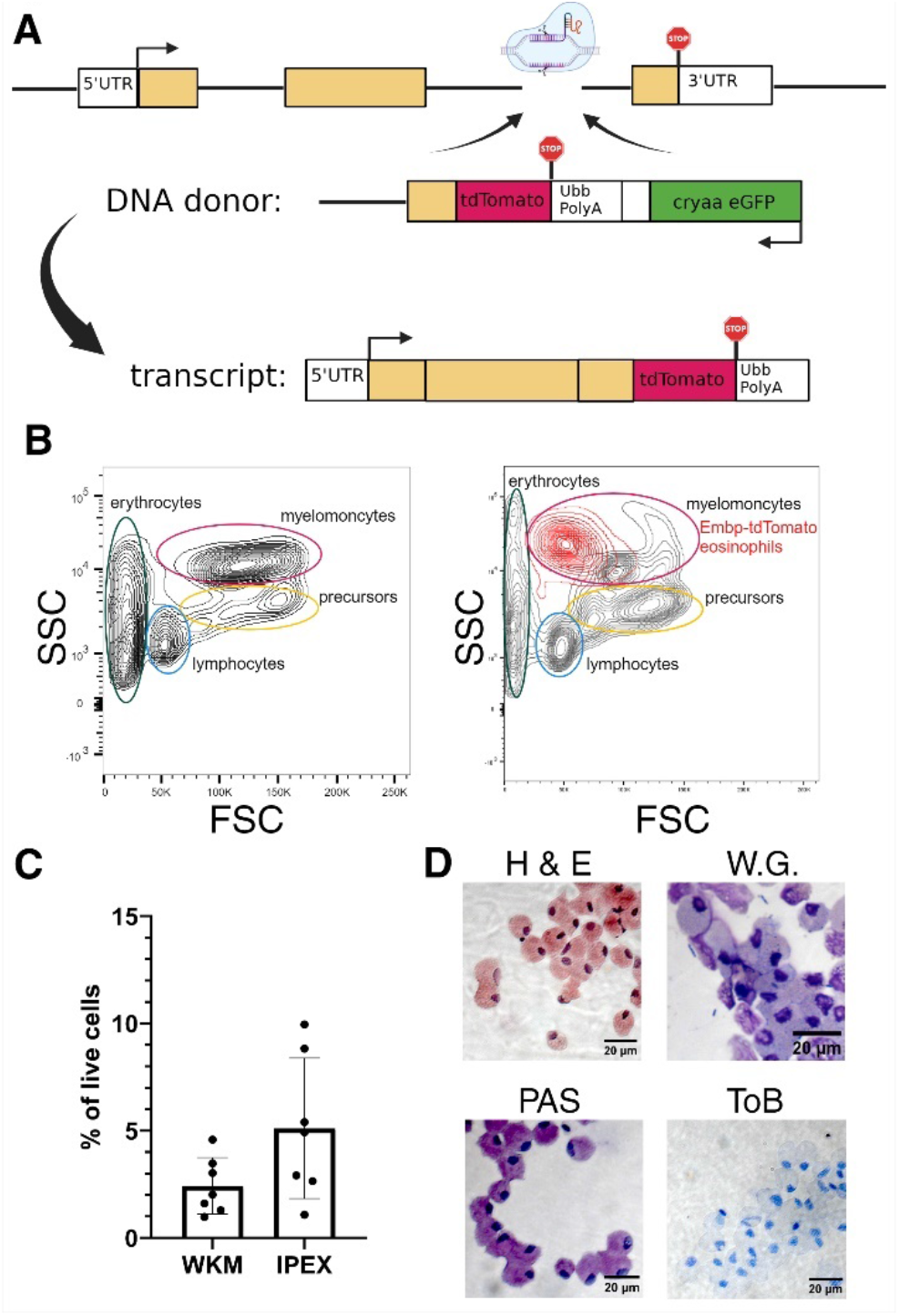
Generation of *TgKI(embp-tdTomato, cryaa:EGFP)* transgenic reporter line and initial validation. (A) Schematic of genetic locus and DNA donor fragment inserted in the *embp* locus. Transcription leads to a fusion construct of Embp and tdTomato. (B) Traditional gating scheme by light-scatter characteristics of a representative wildtype WKM as previously reported^24^. Side by side visualization with the Embp-tdTomato^+^ cell population backgated onto the entire WKM in FSC/SSC. (C) Quantification of Embp-tdTomato^+^ cells in WKM and IPEX samples of this transgenic line, measured by flow cytometry and presentation as proportion of live cells: WKM 2.4% (± 0.38%) and IPEX 5.1% (± 2.7%) mean with SD. (D) FACS sorted Embp-tdTomato^+^ cells stained with Hematoxylin & Eosin (H&E), Wright Giemsa (WG), Periodic Acid Schiff (PAS) and Toluidine Blue (ToB).

Wildtype zebrafish eggs were injected at one cell stage, screened for cryaa:GFP expression at 3 dpf and raised to adulthood. F0 founders were spawned and screened to identify founders transmitting the cryaa:GFP phenotype. cryaa:GFP-positive F1 offspring with stable integration of the transgene were analyzed using flow cytometry for tdTomato expression in the granulocyte compartment. In zebrafish, eosinophils are commonly isolated from two sites: whole kidney marrow (WKM), the bone marrow equivalent of zebrafish, and intraperitoneal exudate (IPEX), cells collected from the peritoneal cavity. tdTomato^+^ cells were identified within both WKM and IPEX populations. These tdTomato-positive cells were brightly fluorescent and appeared within the expected myelomonocytic gate using forward scatter/side scatter gating as has previously been done in zebrafish^24^ (Fig 1B). The proportions of tdTomato^+^ cells in WKM (2.4% ± 0.38%) and IPEX (5.1% ± 2.7%) (Fig 1C) corresponded well to the proportions of eosinophils observed in previous studies with Gata2a^high^ cells. Using FACS, tdTomato^+^ cells were isolated and stained with a panel of histological stains to corroborate the identity of the isolated cells. Using Hematoxylin & Eosin (H&E), Wright Giemsa (WG) and Periodic Acid Schiff (PAS) staining, we found that tdTomato^+^ cells showed the histological hallmarks of zebrafish eosinophils^11^ including a “round to oval shaped nucleus typically located peripherally”^25^ (Fig 1D). We also observed that these cell populations were negative for Toluidine Blue (ToB) staining, which identifies highly granular mast cells which are also known to be present in zebrafish^26^. Confocal imaging of transgenic tdTomato^+^ eosinophils isolated from *TgKI(embp-tdTomato, cryaa:EGFP)*^*ber1*^ animals revealed brightly fluorescent cells with the tagged transgene having a punctate localization that is consistent with targeting of this gene to the granules (Fig 2A). This transgenic reporter could be visualized not only in isolated cell populations, but could also be used to observe eosinophil localization in FFPE sections in conjunction with a commercial mCherry antibody (Fig 2B). Using IF and histological staining of adjacent sections, we found that these tdTomato^+^ transgenic eosinophils localized to the same sites in the zebrafish body as their wildtype counterparts under homeostasis and possessed the histological characteristics of zebrafish eosinophils, suggesting that our tagging approach did not impact eosinophil homeostasis.

**Figure 2.**
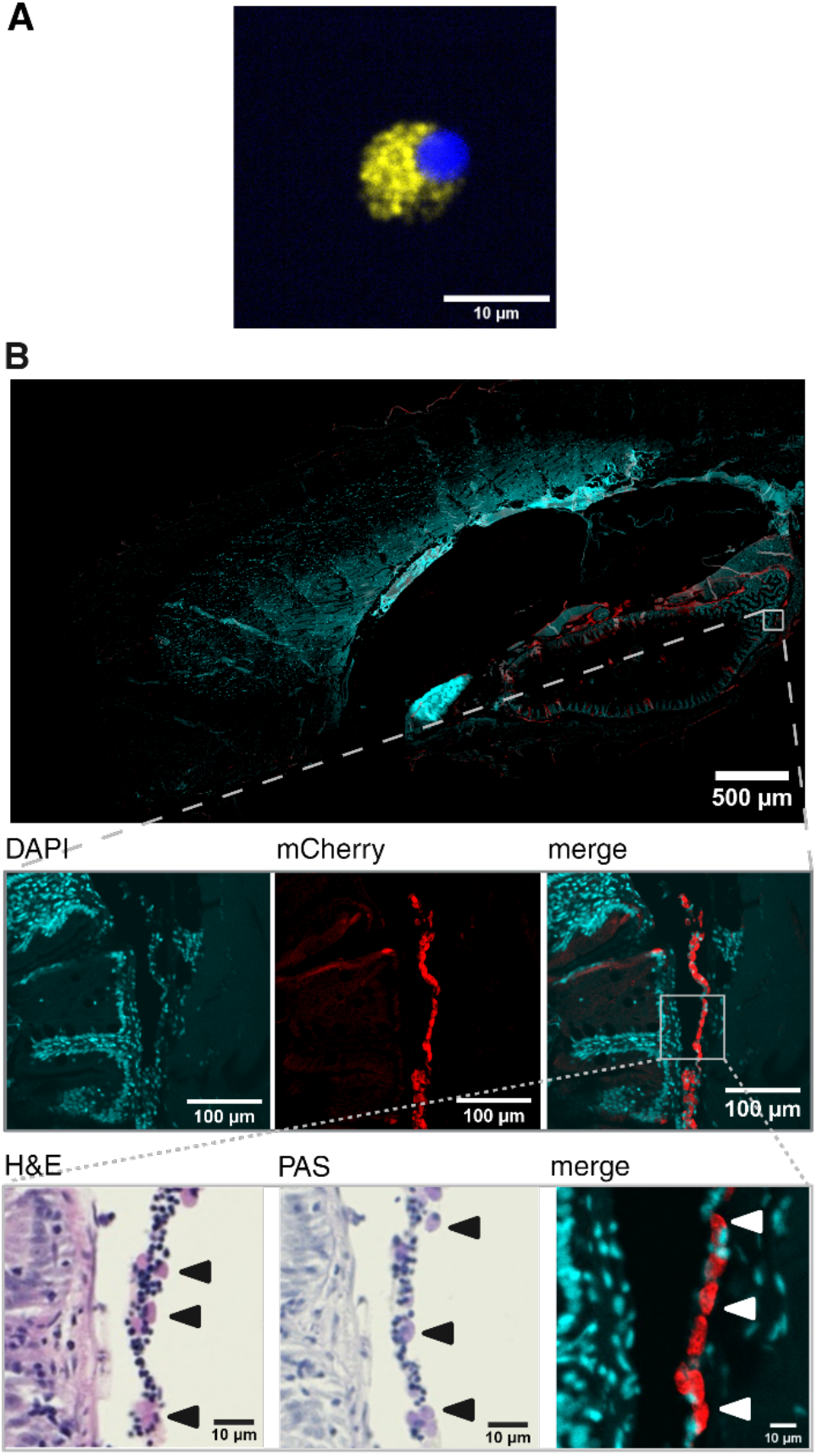
Confocal fluorescent microscopy of *TgKI(embp-tdTomato, cryaa:EGFP)* samples. (A) 40X imaging of a FACS isolated Embp-tdTomato^+^ (yellow) eosinophil. The fluorescent transgene is localized within likely eosinophil granula. Nuclear counterstain with DAPI. (B) Detection of the Embp-tdTomato transgene in 5 μm FFPE sections using mCherry antibodies. Eosinophils are found along the lamina propria lining the intestine and in-between organs, confirmed by H&E and PAS staining of adjacent sections.

In conclusion, endogenous tagging of *embp* with tdTomato created a transgenic reporter line that enabled us to isolation and visualization of zebrafish eosinophils. The transgenically tagged cells showed eosinophil morphology and characteristics by flow cytometry, histological staining and confocal microscopy. Additionally, staining tissue sections with an mCherry antibody confirmed that tdTomato^+^ eosinophils localize like their wildtype counterparts under homeostatic conditions, demonstrating that this targeting approach did not alter eosinophil localization in homeostasis.

Further, using this reporter line, we found that this transgene appeared to localize to granules, suggesting that Embp may be one of the limited subset of eosinophil granular proteins.

Visualizing Embp-tdTomato^+^ eosinophils and wildtype Embp both *in situ* and *ex vivo* While our transgenic reporter line was useful to isolate eosinophils and visualize eosinophils *in vivo*, we also sought to establish a solution to visualize eosinophils in tissue samples that lack the transgenic reporter. To this end, we generated a custom polyclonal antibody targeting Embp (Fig 3A). To confirm eosinophil specificity, the antibody was tested on WKM, IPEX and isolated eosinophils of both *Tg(gata2:eGFP)* and *TgKI(embp-tdTomato, cryaa:EGFP)* lines (Fig 3B+C). Positive staining was observed in 92.45%/87.47% (Embp/Gata2a line) of sorted eosinophils, 4.62%/3.58% (Embp/Gata2a line) in WKM and 12.63%/9.86% (Embp/Gata2a line) in IPEX samples, consistent with previous reports on the proportion of eosinophils in such samples. Importantly, staining of GFP^-^ and tdTomato^-^ fractions from *Tg(Gata2a:GFP)* or *TgKI(embp-tdTomato, cryaa:EGFP)* revealed no staining in eosinophil-depleted fractions by the antibody highlighting its specificity (Fig S1B).

**Figure 3.**
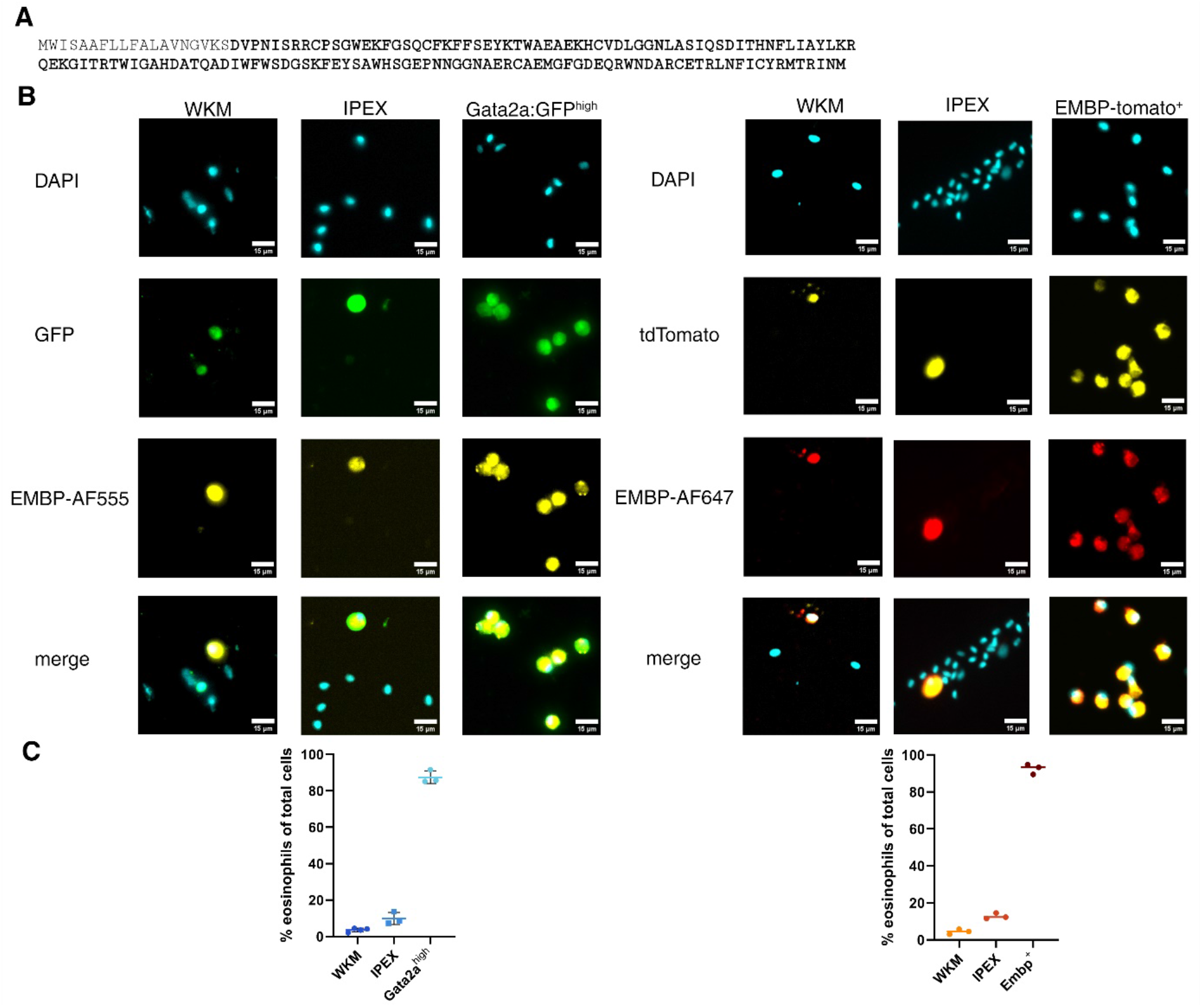
Novel polyclonal antibody targeting Embp using in immunofluorescence staining. (A) Amino acid sequence of Embp in single letter code. In bold: the part of the antigen used to generate the polyclonal antibody. (B) Staining of isolated WKM, IPEX and eosinophil cell populations of both the *Tg(gata2:eGFP)* and *TgKI(embp-tdTomato,cryaa:EGFP)* lines. Eosinophils were sorted by transgenic reporter expression (either Gata2a^high^ or Embp^+^) (C) Quantification of cells stained positively for Embp with polyclonal antibody in the afore mentioned sorted cell populations. Positive staining was assessed in a minimum of six random fields of view across three independent experiments. 300-500 cells were evaluated per sample. Data represented as percentage of positive Embp staining in total cell population, mean with SD of each independent experiment.

We also sought to determine whether this antibody could also be used to identify eosinophils in paraffin embedded tissues. Antibody staining of formalin-fixed paraffin-embedded (FFPE) tissue sections (Fig 4) demonstrated that this antibody could detect eosinophils within the tissue, with the staining patterns corresponding with the expected distribution of eosinophils within the tissue. To further corroborate stained cells as eosinophils, we stained adjacent sections with H&E and PAS (Fig 4A) confirming that cells positive for Embp staining localized to the same locations as eosinophils detected by H&E and PAS. Eosinophils were found mainly in clusters in between organs of the peritoneal cavity, along the basal lamina propria in the gut and intestine as well as in the WKM. Negligible antibody staining was observed in regions of the animal where few if any eosinophils were observed, confirming the specificity of the antibody binding and efficacy of this reagent for the detection of endogenous eosinophils.

**Figure 4.**
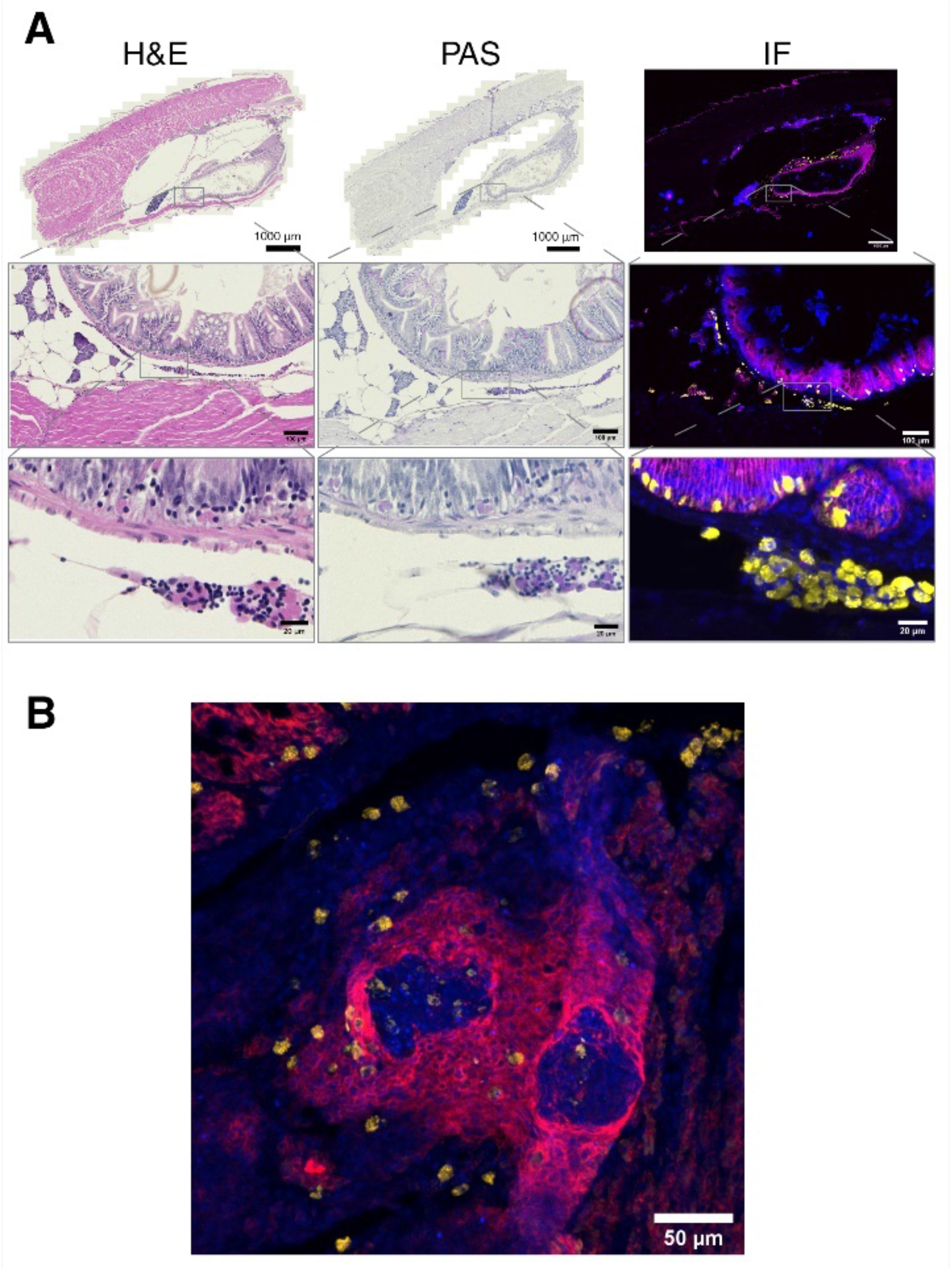
Embp antibody staining in FFPE tissue samples. (A) H&E, PAS and IF staining with Embp antibody in healthy *TgKI(embp-tdTomato,cryaa:EGFP)* adults. Blue: DAPI Yellow: eosinophils Magenta: E-Cadherin staining epithelial cells. (B) Staining of eosinophils in *M. marinum* infected zebrafish FFPE sections. Eosinophils can be found in physiological sites in the body as well as in granuloma. Magenta: E-cadherin, marking epithelioid macrophages surrounding the necrotic core of the granuloma. Yellow: eosinophils stained by Embp antibody. Blue: DAPI.

During inflammation, immune populations including eosinophils undergo profound transcriptional changes and eosinophils can also undergo degranulation, processes that can shift reporter gene expression and localization. To confirm that our antibody and the broader reporter approach could still be used to detect eosinophils within inflamed tissues, we visualized eosinophil localization in *Mycobacterium marinum* infected FFPE samples of adult zebrafish (Fig 4B). *M. marinum* infection causes granulomatous inflammation in zebrafish and eosinophils have been found within the zebrafish granuloma^27^. We found that eosinophils could be stained within the mycobacterial granuloma as well as in surrounding tissue indicating that even under inflamed conditions, this marker could be used to identify eosinophils. This finding is also consistent with previous studies demonstrating recruitment of eosinophils to mycobacterial granulomas, providing further evidence for eosinophils within this structure and the spatial distribution of the cells throughout the granuloma.

Our previous experiments have indicated that not only is Embp an effective marker of eosinophils, but have also supported a putative eosinophil granular localization for Embp. In mammals, eosinophil granules contain a limited array of highly expressed effector genes. To definitively establish Embp as a granular protein we employed cryo-immunogold electron microscopy (EM) to localize Embp within the eosinophil. Embp-tdTomato^+^ eosinophils were isolated from *TgKI(embp-tdTomato, cryaa:EGFP)* animals by flow cytometry and stained with both our polyclonal Embp antibody to detect the native antigen and an mCherry antibody, to visualize transgene localization with the same cells. We found that both the endogenous Embp and the tdTomato transgene are almost entirely localized within the granules of the eosinophils, with only sporadic protein detection in the cytoplasm (Fig 5). Thus our findings establish Embp as a *bona fide* eosinophil granular protein in zebrafish, and a potential role for this protein as an eosinophil effector protein.

**Figure 5.**
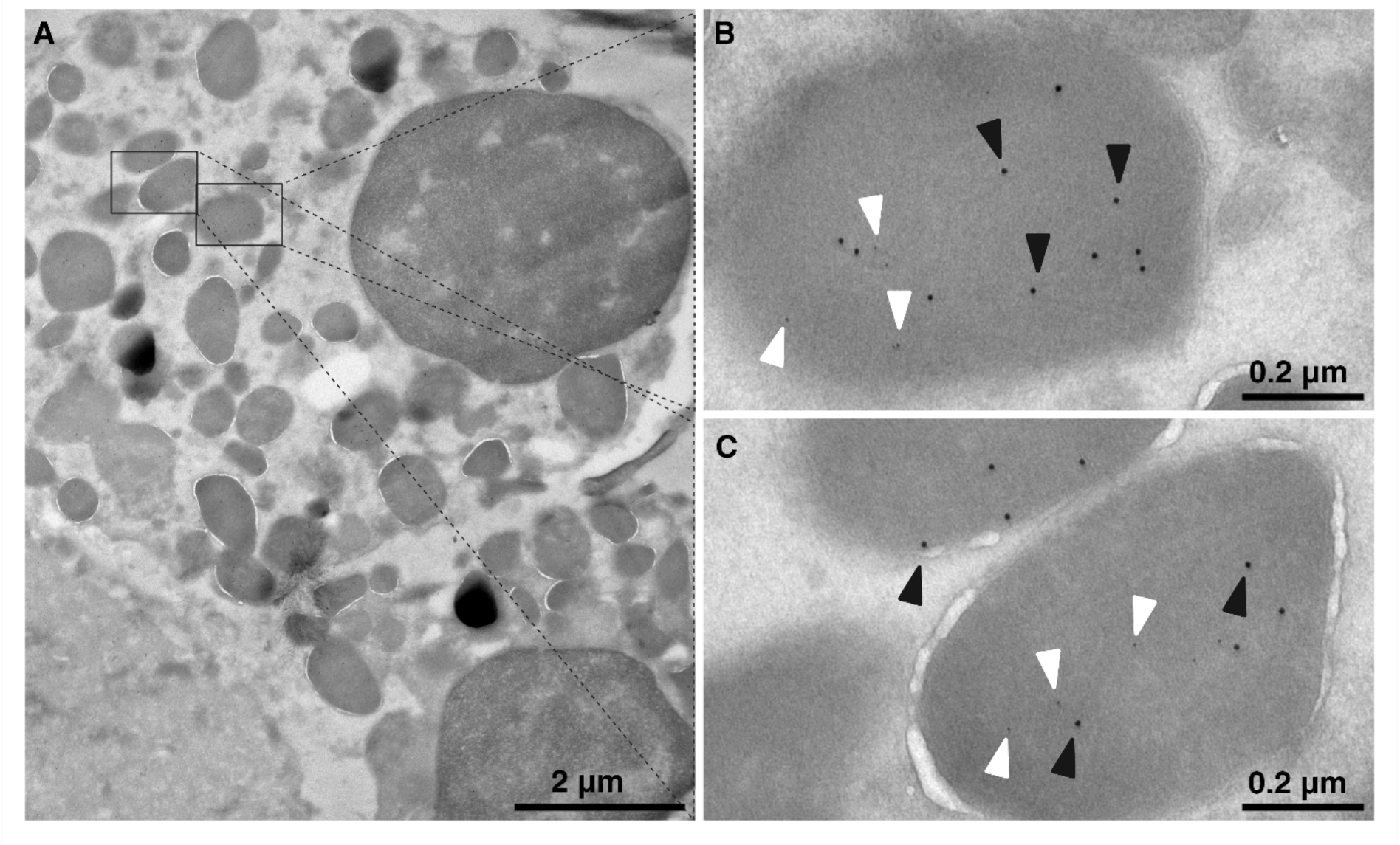
Cryo-immunogold EM of FACS isolated Embp-tdTomato^+^ eosinophils. (A) Overview of an eosinophil. (B+C) Zoom in on two different granula of this cell. Embp antibody is visualized by a 12 nm gold secondary antibody (black arrow heads) and mCherry antibody is visualized by 6 nm gold secondary antibody (white arrow heads). Both target antigens detected in the same granule.

## Discussion

Eosinophils are critical effectors of the vertebrate immune response, contributing to the control of helminth pathogens and tissue remodeling but also leading to detrimental pathology in diseases such as asthma^28^. Eosinophils are broadly conserved in vertebrates, but despite their recognition in zebrafish^25^, tools to study this cell population and their specialized effectors are lacking in this emerging infection model^29^. In this study, we identified a novel highly specific marker for zebrafish eosinophil granules, Embp. We initially identified Embp from analysis of previously published RNA-Seq data sets, which suggested that *embp* (also known as *dkeyp75b4*.*10* ENSDART00000143719, ENSDART00000110749) was both highly and differentially expressed in eosinophilic granulocytes, with minimal expression in other hematopoietic-derived lineages. This eosinophil-specific expression of Embp is consistent with recent observations of this gene and its promoter^23^, though the localization and function of this gene was unclear. We generated a transgenic reporter line tagging the endogenous Embp with a fluorescent protein on its 3’ end by CRISPR insertion to investigate the localization of this gene resulting in the endogenous promoter of *embp* driving the expression of a full-length transgenic Embp-tdTomato construct. Using this zebrafish line, we found that *embp* was specifically expressed within eosinophils and not in other immune populations, demonstrating that this line could be used to specifically mark eosinophils in zebrafish. This marker line could be used for flow cytometry, confocal microscopy and histochemistry to identity eosinophils, expanding the toolkit available to study the function of these cells in zebrafish. In addition, a newly generated polyclonal antibody targeting Embp allows visualization of endogenous eosinophils *in vivo* and *ex vivo*.

Comparing the proportions of eosinophils in the previously characterized *Tg(gata2:eGFP)* line and our new *TgKI(embp-tdTomato, cryaa:EGFP)* line, we find very similar levels of our cells of interest. While numbers were close between the two lines, we did find a slight statistically significant increase in the number of eosinophils only in the WKM population of *TgKI(embp-tdTomato, cryaa:EGFP)* (Fig S1) but not in IPEX populations. Considering the low number of eosinophils in WKM and the high variability in eosinophil number per animal in our data and the data of other groups^25^, we expect that this could potentially be driven by stochastic differences in this small group of animals. A second possibility is that the distinct fluorophores and fluorescent intensity of each reporter could lead to subtle differences in the detection frequency. Overall, however, the relative number of eosinophils in the new *TgKI(embp-tdTomato, cryaa:EGFP)* line falls within the range of previously reported eosinophil counts^11^.

Structurally Embp possesses a c-type lectin domain. Although there are a number of C-type lectin proteins expressed in mammalian eosinophils including major basic protein 1 (MBP-1)^30^, major basic protein 2 (MBP-2)^31^, Dectin-1^32^ and C-type lectin domain family 4, member a4 (Clec4a4)^33^, MBP-1 and MBP-2 are the only C-type lectins that are canonical granule proteins in mammalian eosinophils^8^. Our cryo-EM data (Fig 5) together with confocal microscopy (Fig 2), demonstrate that in zebrafish Embp is an eosinophil granule protein, likely acting as an eosinophil effector molecule. MBP-1 and MBP-2 are amongst the most highly expressed proteins in mammalian eosinophils. Similarly, *embp* was identified as one of the most highly expressed genes in previous scRNA-Seq data and both our reporter and antibody show robust positivity indicating high level expression. Based on the strong expression of Embp and localization within the granules, its exclusivity to eosinophils and its C-type lectin domain, we expect that Embp is a zebrafish ortholog of the MBP family. Thus, in light of our data, we believe it is accurate to name this gene Embp rather than the previous gene name, *dkeyp75b4*.*10*, or the recently proposed eslec.

In mammals, eosinophil major basic protein 1 is, as the name suggests, characterized by a strongly basic pI. Surprisingly, the pI of Embp is relatively neutral (pI 6.8). However, while hMBP-1 is highly basic (pI 11.4), hMBP-2 is known to also have a pI much closer to neutral (pI 8.7). As we saw in zebrafish for Embp, hMBP-2 is exclusively expressed in eosinophils^34^, while hMBP-1, although predominantly expressed in eosinophils is also expressed in basophils and mast cells, suggesting that *embp* may be the ortholog of hMBP-2. Interestingly, in ruminants, MBP-2 is the only MBP present^35^. Also, the relatively neutral pI of Embp suggests that the characteristically strongly basic pI of hMBP-1 may be a relatively recent evolutionary occurance. Indeed, the pIs of MBP orthologues vary considerably across species^34,35^. Although further studies into the biochemical and functional properties of this zebrafish orthologue will be needed, zebrafish *embp* will likely offer an interesting window into the evolution of MBP proteins.

Addressing a critical gap in the study of zebrafish eosinophils, the new tools presented here will also allow to study the development and maturation of eosinophils as well as function of eosinophils in diverse diseases. Recent efforts using zebrafish to model allergic inflammation and food allergies have identified granulocytes as significant contributors to these conditions^36^, but they have not been able to identify the specific cell types responsible. By facilitating studies of eosinophil function in zebrafish, our tools will enable researchers to discover new roles for eosinophils that can be conserved in human disease. As with the MBP protein family, the similarities and differences in eosinophil responses in zebrafish will also offer a window into the evolution of these cells, as zebrafish are one of the most distant human ancestors to have eosinophils. Use of zebrafish for comparative immunology approaches will highlight centrally conserved immune responses and eosinophil behavior that function across vertebrate species.

## Methods

### Zebrafish housing and breeding

Animals were housed and bred under standard conditions. All experiments were conducted in accordance with institutional (Max Planck Institute for Infection Biology), state (LAGeSo Berlin), and German ethical and animal welfare guidelines and regulations. Male and female animals were used in approximately the same ratios for all experiments.

### Zebrafish lines

The wild-type strain in this study was *AB and all zebrafish strains used this background. Transgenic lines used in this study: *Tg(gata2:eGFP)* that has been previously been described^12^ and the newly created *TgKI(embp-tdTomato,cryaa:EGFP)*^*ber1*^ line (see below). Fluorescent reporter lines were maintained as heterozygotes.

### Generation of transgenic zebrafish line

The new eosinophil reporter line was created using a CRISPR Cas9 based knock-in approach inserting into a non-coding region as previously described^13^. The transgenic insert used to generate *TgKI(embp-tdTomato, cryaa:EGFP)* zebrafish was assembled using the Golden Gate cloning system^14^. The insert combines the last two exons of *embp* (also known as *dkeyp75b4*.*10* or *eslec*, ENSDARG00000079043) amplified from bacterial artificial chromosome (BAC) clone CH211-226G7 (BACPAC Genomics), a tdTomato sequence followed by UbB polyA and a transgenesis marker encompassing the *cryaa* promoter^15^ driving EGFP in the reverse orientation. The DNA donor fragment was PCR amplified and cleaned up prior to injection using the Gel & PCR Clean-Up Kit (Macherey-Nagel). One cell stage zebrafish embryos were injected with ∼1-2 nl of 2.5 ng/μl DNA donor fragment together with 12.5 ng/μl sgRNA (GGACAATTTTCATGACAAGA, ordered from IDT) and 75 ng/μl Cas9 protein (IDT). 3 dpf larva were screened for GFP expression in the retina and raised to adult hood. Mosaic F0 founders were outcrossed to *AB fish and screened for germline transmission of the transgenesis marker (GFP^+^ eyes). GFP^+^ F1 fish were raised to adulthood and screened for tdTomato^+^ eosinophils using flow cytometry.

### Flow cytometry and cell sorting

Immune cells were isolated from whole kidney marrow (WKM) and intraperitoneal exudate (IPEX) in L-15 (Thermo Fisher Scientific) + 5% FCS as previously described^11^. In brief, an incision was made on the ventral side of the fish and the peritoneal cavity washed with 100 μl L-15 + 5% FCS to collect IPEX cells. This process was repeated ∼3-4 times to collect a sufficient number of cells. Next, the inner organs were removed and WKM isolated as previously described^16^. Single cell suspensions were made using 100 μm and 40 μm cell strainers (VWR). The single cell suspensions were centrifuged at 300xg for 5 min at room temperature (RT) and resuspended in 1 ml 1X PBS + 5% FCS. Cells were stained with DAPI (Thermo Fisher Scientific) for 15 min at RT before flow cytometric analysis. Flow cytometry was performed on a BD FACS Canto II (Becton Dickinson) and cell sorting was performed on a Sony MA900 cell sorter (Sony). Data were analyzed with FlowJo version 10.6.2.

### Cytology

Cytospin preparations were carried out using a Cytospin3 (Shandon) centrifuge. Sorted eosinophils were resuspended in 150 μl of 1X PBS with 5% FCS. Cytospin chambers were pre-wetted with 100 μl of 5% FCS-PBS. The cells were then centrifuged at 800 rpm for 3 min at low acceleration. Slides were air-dried overnight at RT before fixation with 4% paraformaldehyde (PFA) in PBS and subsequent staining.

### Zebrafish sectioning

Zebrafish were euthanized by hypothermic shock in ice cold water. Subsequently, the fish were fixed in 4% PFA solution for 2 days at RT, decalcified in 0.35 M EDTA solution for 7 days and paraffin embedded as previously described^17^. Paraffin embedded zebrafish were sectioned into 5 μm thin sections using a Microm HM355S Rotary Microtome (Thermo Fisher Scientific), floated onto polysine slides (VWR) and dried at 37°C overnight.

### Histological staining

Paraffin sections were deparaffinized in two changes of 100% xylene for 10 min each and then rehydrated in two changes of 100% EtOH, two changes of 95% EtOH and one change each of 70% EtOH and 50% EtOH. Sections were washed with deionized water.

Histological staining was carried out according to the manufacturer’s protocols (Sigma-Aldrich). Samples were mounted using DPX new mounting media (Sigma-Aldrich).

### Immunofluorescent (IF) Staining

Single cell suspensions were obtained as described in the “Flow cytometry and cell sorting” section. Tissue sections were obtained as described under “Zebrafish sectioning” and deparaffinized as described in “Histological staining”. The *M. marinum* infected, paraffin-embedded zebrafish were a gift from David M. Tobin.

Single cell suspensions were fixed with 4% PFA in PBS in a 96 well plate for 15 min at RT. After washing 3x with 1X PBS, cells were permeabilized with 0.1% (v/v) Triton-X/PBS for 15 min at. Blocking was done with 3% goat serum/PBS for 1 h at RT and subsequent primary antibody (Table S1) staining was done in 3% goat serum/PBS overnight at 4°C. Cells were washed 3x with 1X PBS and stained with secondary antibody (Table S1) solution in 3% goat serum/PBS for 1 h at RT. Cells were washed again 3x with 1X PBS and counterstained with DAPI (0.2 μg/mL). After a final wash with 1X PBS, cells were transferred to 50% (v/v) glycerol in 20 mM TRIS mounting media for subsequent imaging.

For histology slides, antigen retrieval was performed by incubating deparaffinized slide in antigen retrieval solution of 10 mM Tris/1 mM EDTA, pH 9.0 followed by heat treatment in a pressure cooker (Krups) for 10 min. After cooling down to RT in antigen retrieval buffer, slides were blocked with 3% goat serum/PBS for min. 30 min and subsequently stained as described above. Selected samples were additionally treated with Vector TrueVIEW Autofluorescence Quenching Kit (Vector Laboratories) according to the manufacturer’s protocol and mounted using the hard-curing mounting medium provided with the kit.

### Microscopy

Color images of classical histological staining were acquired using a Leica DMR microscope and ProgRes SpeedX core3 camera (Jenoptik) or AxioScan Z1 (Zeiss).

Fluorescence images were taken on a Nikon ECLIPSE Ti2 microscope (Nikon) with a Dragonfly 200 confocal system (Oxford Instruments) and 10X, 20X or 40X objectives (Nikon). Image files were processed with Fiji ImageJ^18^ and Zeiss Zen Blue (Zeiss).

### Cryo-Immunogold Electron Microscopy (EM)

Embp-tdTomato^+^ eosinophils were isolated and sorted as described above. Cells were fixed with 4% PFA and 0.05% GA in PBS. The cells were harvested, gelatin-embedded and infiltrated with 2.3 M Sucrose according to the method described^19^. Ultrathin sections were cut at -110°C with a RMC MTX/CRX cryo-ultramicrotome (Boeckeler Instruments Inc., Tucson AZ, USA) transferred to carbon- and pioloform-coated EM-grids and blocked with 0.3% BSA, 6 mM Glycin, 3% CWFG in PBS. The sections were incubated with appropriate dilutions in the same buffer of rabbit polyclonal antibody directed against Embp (Table S1) and chicken monoclonal antibody against mCherry (Table S1). Secondary antibody-incubations were carried out with goat-anti-rabbit or anti-chicken antibodies coupled to 12 or 6 nm gold particles, respectively (Jackson Immuno Research, West Grove, PA, USA). Specimens were then contrasted and embedded with uranyl-acetate/methyl-cellulose following the method described before^20^ and analyzed in a Leo 912AB transmission electron microscope (Zeiss, Oberkochen, Germany) at 120 kV acceleration voltage. Micrograph-mosaics were scanned using a bottom mount Cantega digital camera (SIS, Münster, Germany) with ImageSP software from TRS (Tröndle, Moorenweis, Germany). Images were processed with FigureJ^21^ in Fiji ImageJ.

### Statistics

All experiments were repeated at least three times independently. Group sizes and statistical tests used are indicated in the figures and figure legends. Statistical tests were performed and visualized using Prism 9 (GraphPad).

## Acknowledgments

This work was supported by the Max Planck Society. In addition, MH is supported by the Add-On Fellowship of the Joachim Herz Foundation. We would like to thank the Zychlinsky department for sharing lab equipment and reagents as well as helpful discussion. We would like to thank Charlie J. Pyle and David M. Tobin for sharing material and Silke Lehmann, Marten Walk, Carsten Weiland, Janine Bleske, Jens Otto and Ines Neumann for excellent care of the zebrafish facility. Figures were created with BioRender.com.

## Authorship Contributions

MH designed research, performed research, collected data, analyzed and interpreted data, performed statistical analysis, wrote the manuscript. CG performed research, collected data, analyzed and interpreted data. VB performed research, collected data, analyzed and interpreted data. CD performed research and collected data. MRC designed research, performed research, collected data, analyzed and interpreted data, wrote the manuscript and supervision.

