## Supplementary figures and tables for "Identification of a specific granular marker of zebrafish eosinophils enables development of new tools for their study"

*Table S1 Antibodies used in this study*

| ANTIBODY | SOURCE | IDENTIFIER |
|---|---|---|
| **E-CADHERIN CLONE 36** | BD Transduction Laboratories | 610181 |
| **DKEYP75B4.10** | Boster Bio | DZ41383 |
| **GFP** | Aves Labs | GFP-1020 |
| **MCHERRY** | Aves Labs | MCHERRY-0100 |
| **ALEXAFLUOR 647 GOAT ANTI-MOUSE** | Thermo Fisher | A-21235 |
| **ALEXAFLUOR 555 GOAT ANTI-RABBIT** | Invitrogen | A32732 |
| **ALEXAFLUOR 647 GOAT ANTI-RABBIT** | Invitrogen | A32733 |
| **ALEXAFLUOR 488 GOAT ANTI-CHICKEN IGY** | Thermo Fisher | A-11039 |
| **ALEXAFLUOR 647 GOAT ANTI-CHICKEN IGY** | Thermo Fisher | A32933 |
| **6 NM COLLOIDAL GOLD GOAT ANTI-CHICKEN IGY** | Jackson Immuno Research | 103-005-155 |
| **12 NM COLLOIDAL GOLD GOAT ANTI-RABBIT** | Jackson Immuno Research | 111-205-144 |

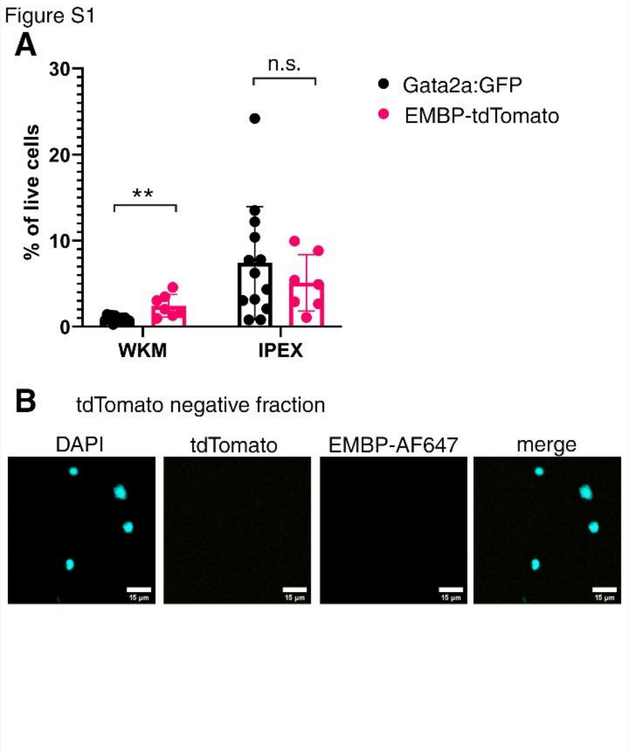


Figure S1. (A) Comparison of the relative abundance of eosinophils in flow cytometry of WKM and IPEX between the *Tg(gata2:eGFP)* and *TgKI(embp-tdTomato,cryaa:EGFP)* lines. Data represented as mean percentage of live cells with SD. Students t-test, ** $p < 0.01$ with 95% CI (B) tdTomato$^{-}$ cells sorted from from *TgKI(embp-tdTomato, cryaa:EGFP)* zebrafish line. Stained with Embp antibody and DAPI.
